# Practical Guidelines for Secure Cloud Computing for Genomic Data

**DOI:** 10.1101/034876

**Authors:** Somalee Datta, Keith Bettinger, Michael Snyder

## Cloud security challenges

Large scale genomics studies involving thousands of whole genome or exome sequences are underway^1^ on Cloud. While Cloud provides many conveniences for genomics research, it also raises concerns regarding large scale hacking, bad press, potential loss of patient privacy and the resulting loss of patient trust. Cloud providers argue that they have significant investments and expertise in security and, therefore, Cloud is equally secure, if not more so, compared to on-premise infrastructure. This gap in assessment of Cloud security is, in part, due to a fast evolving and largely unfamiliar technology stack for genomics data owners.

What makes the Cloud security landscape discussion challenging is that security recommendations differ across regulatory bodies, besides being inconsistent between on-premise and Cloud requirements. For example, Institutional Review Board (IRB) often require Health Insurance Portability and Accountability Act^2^ (HIPAA) level Cloud security even for non Protected Health Information^3^ (PHI) data. In another example, Database of Genotypes and Phenotypes^4^ (dbGAP) has different encryption requirements for on-premise and Cloud environments.

Our intent in this Commentary is to provide the genomics community with a set of Cloud security guidelines that will meet a wide range of regulatory requirements. Although the Cloud technology stack will continue to evolve rapidly, thus changing the specifics of implementation, we believe that these guidelines will be applicable for the foreseeable future.

## Cloud security guidelines

At Stanford, we have developed a secure Cloud gateway to support multiple projects including NIH and other government agency datasets as well as private datasets. This gateway supports multiple regulatory requirements from various agencies. We highlight these requirements and make recommendations for institutional Cloud administrators to implement. We will specifically discuss requirements for non-PHI data covered under a typical genomic Data Use Agreement (DUA) but our guidelines incorporate HIPAA requirements. We will present specific discussion for Google Cloud Platform (GCP), based on our own integration experience, in order to illustrate some of the requirements, but the principle behind the implementation is applicable on all Clouds.

Note that Genomics Cloud Service providers such as DNAnexus or Seven Bridges Genomics must be held to the same standards of security as the underlying Cloud infrastructure-as-a-service^5^ provider such as Amazon AWS or GCP or Microsoft Azure. Figure 1 shows the relationship between the underlying Cloud infrastructure security features and institutional administrative effort needed to make the Cloud secure for researchers. In our administrative experience, ready-to-use features, presence of Cloud APIs, clarity of documentation and quick support turnaround make an administrator’s task significantly easier. It is our belief that without institutional administrator efforts, no Cloud infrastructure-as-a-service can meet security requirements out-of-the-box.

**Figure 1:**
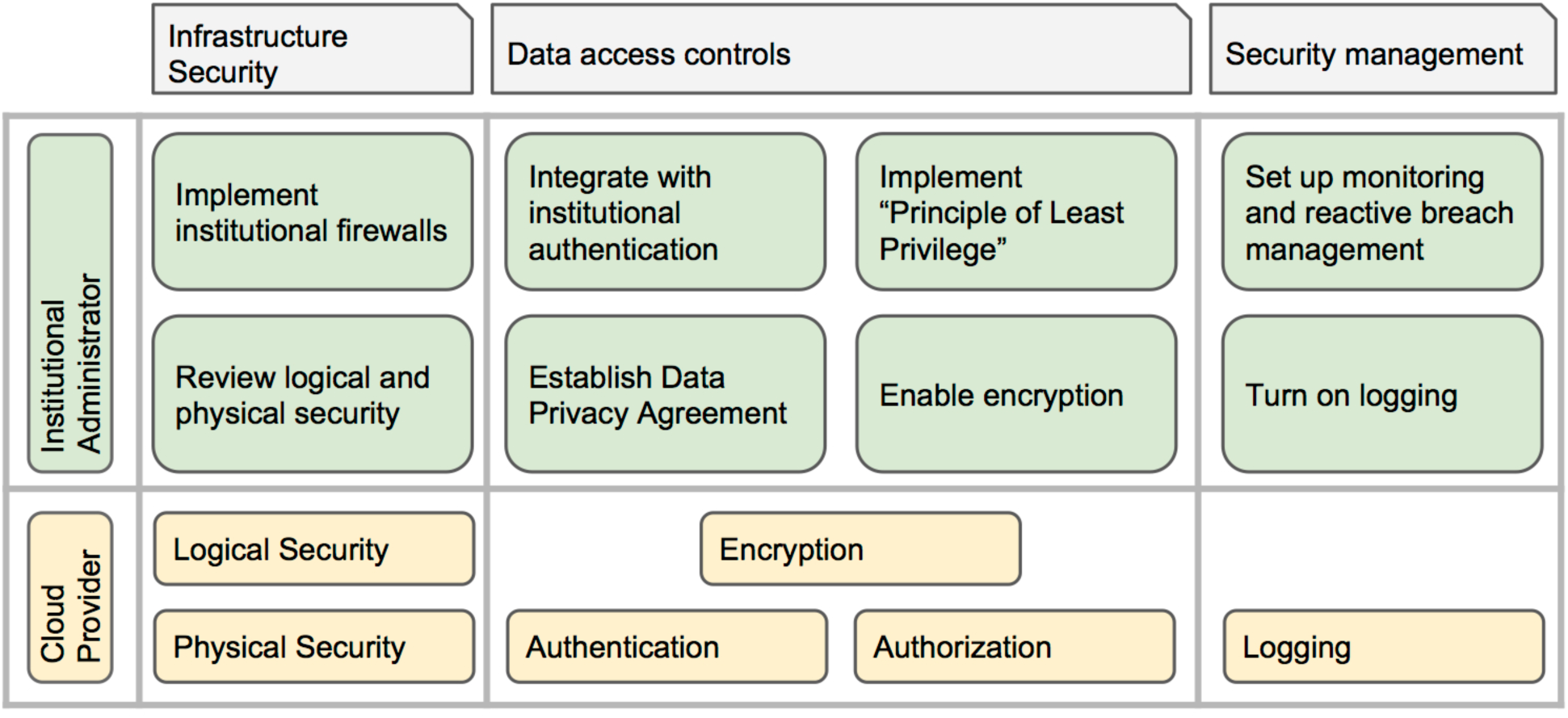
This figure shows the high level features in light grey boxes - infrastructure security, data access controls and security management - that cloud security must meet for genomics research. The boxes at the bottom in yellow background present the Cloud infrastructure-as-a-service capabilities. The light green boxes in the middle present institutional administrator responsibilities.

**Data Privacy Agreement**: Data privacy guidelines impact non-public data. The extent of impact depends on regulations around the specific data set. It is recommended that before transferring genomic data to the Cloud, you work with your Information Officer or legal division to set up a Service Agreement with the Cloud provider that meets Institutional data use and privacy policies.

**Physical and Logical Security:** In physical security, Cloud providers must follow best practices in Data Center access control systems, alarms systems, hardware tracking and disposal. In logical security, Cloud providers must follow best practices regarding malware monitoring and prevention, vulnerability identification and remediation.

Physical and logical security requirements are data dependent. For example, for dbGaP and HIPAA compliance, it is required that hard drives be destroyed before disposal *e.g*. before returning bad disks to vendor as part of service contract. Some studies specifically require the Cloud to have HIPAA compliance or Federal Information Security Management Act^6^ (FISMA) compliance or Federal Risk and Authorization Management Program^7^ (FedRAMP) compliance. We believe that in time, all major Cloud providers will support FedRAMP.

A Cloud provider will typically have regular third party audits for compliance standards. Administrators should review these certifications annually against institutional and project requirements. GCP, as illustration, has regular third party audits^8^ for SOC2 / SOC3^9^ and ISO27001^9^ standards.

**Encryption**: Typically, encryption at rest and in transit is only required for PHI data. However, dbGaP requires encryption on Cloud (and not on-premise) and we are increasingly seeing this requirement in IRB guided studies.

We caution the user that not all Cloud providers have server side encryption^10^ as a service and therefore, the administrator must pay particular attention to feature availability and roadmap. For example, Google Cloud encrypts all data by default before it is written to disk and is encrypted in transit between services. In this process, Google manages the cryptographic keys on the user’s behalf using the same hardened key management systems that Google uses for its own encrypted data, including strict key access controls and auditing. Each Cloud Storage object’s data and metadata is encrypted under the 128-bit Advanced Encryption Standard^11^ (AES-128), and each encryption key is itself encrypted with a regularly rotated set of master keys.

When server side encryption is not available, users become responsible for client side encryption and this responsibility includes complex key management process. We believe this to be undue risk for administrators without appropriate IT software support.

**Authentication:** An authentication process confirms user’s identity and is used to manage access to data. This is required for any non-public data. At Stanford, when the researcher is no longer affiliated with the University, they lose their institutional ID and, consequently, they lose access to Stanford managed systems. We recommend integration of the Cloud with the institute’s authentication system so that institution retains control on data access. Such integration allows for institutional policies to be followed regarding training requirements. Additionally, such integration may provide some protection against hacking attacks by relying on institution’s ability to identify and react to such attacks.

We realize that the above recommendation poses challenge for the researcher in the scenario where the researcher leaves the institute but retains access to data. In this case, one option would be to retain access to the institute’s identity for the duration of the project. Another option would be to move (or copy) the data to the new location. Choice of a specific option should be guided on a case-by-case basis and include consideration towards complexity of data movement and remaining duration of the project. If, on the other hand, researcher loses access to the data but retains institutional identity, the burden is on Principal Investigator (PI) to inform the administrator who will need to explicitly remove access.

Not all Cloud providers support institutional integration. If a Cloud provider does not provide such mechanism, administrator will specifically need to understand provider’s ability to securely manage logins and passwords. Additionally, the administrator will need to take on the risk of managing account activations and deactivations.

**Principle of Least Privilege (PLP):** PLP means that any user has the minimum number of privileges necessary to accomplish the work done in their role in the system. This requirement is typically true for HIPAA compliance. PLP can significantly reduce risk of accidental data exposure.

To illustrate PLP, consider the example where two researchers, one bioinformatician and one geneticist are granted access to data from a DNA-Seq study. With PLP in place, the bioinformatician will have access to the raw sequence data, aligned sequence data, variant calls and annotated variant calls. Only the bioinformatician will be able to run variant calling and annotation pipelines on the sequence data to generate variant calls and annotated variant calls. The geneticist will have access to aligned sequence data, annotated variant calls, phenotypic data and interpretation report. And only the geneticist will be able to create an interpreted report.

In order to support PLP, the Cloud provider must support fine grain management of Access Control List (ACL), preferably independently on compute and storage units. We find that while it is technically trivial to set up a PLP, the change management of PLP over the lifetime of a project needs the administrator to be closely familiar with the research workflow to support ongoing changes in ACLs. A separate and auditable change control process is recommended.

**Firewalls:** The network security of infrastructure is one of the biggest concerns around use of Cloud for genomic data. We found significant differences between Cloud providers in how the network infrastructure is managed and we recommend that Cloud administrator pay particular attention to how network restriction will be implemented.

In the case of GCP, access methods like the Secure Shell (SSH) service, are allowed to originate from anywhere in the world thus allowing researcher to connect to their Virtual Machines (VMs) from anywhere in the world. Unfortunately, robotic hacking attempts also use the same access mechanism. VM will deny the connection to hacking machine due to failure of authentication but in doing so, a few kilobyte egress occurs. These egress are logged as billing events and persistent hacking attempts over period of week can show up as a few megabytes of egress activity.

In absence of firewall restrictions, Cloud framework, in order to provide better response to user, can automatically migrate the VM and related data to a Data Center near Internet Protocol (IP) address origin. This can potentially cause DUA violations if data leaves national boundary.

To prevent these types of scenarios, firewalls for VMs should be configured to only allow SSH and HyperText Transfer Protocol over Secure Socket Layer (HTTPS) access from a restricted set of source IP addresses. This set should be no larger than the set of addresses at the institution, and ideally would be restricted to the particular IP addresses associated with the researchers involved in the Cloud-based analysis.

In addition, we also found the need for the firewall itself to be configured to ignore rather than reject filtered traffic -- the difference being that rejecting a connection attempt itself generates data traffic when the VM replies to the connecting machine, whereas ignoring a connection attempt does not generate any additional data traffic.

We also observed that many VM images within the GCP try to update their Operating System (OS) packages automatically, some as often as nightly. For some OSes *(e.g*., CentOS), the update procedure accesses a main repository for a list of machines from which an update can be obtained. This list of machines can include IP addresses from geographical locations around the world, and the update procedure can select from any of them. These updates can then generate data accesses for regions outside of the U.S., which may pose audit risk as well. Administrators can provide startup script to deactivate this updating procedure, preventing any such data accesses from showing up in the logs. Preventing updates during the running of a VM is also a good practice independent of the data access issue, to maintain control of the exact software environment under which an analysis is run.

In summary, after network firewall implementation, it is best to test for unexpected egress or data access patterns.

**Logging & Monitoring:** With any secure system, administrators assume that mistakes can happen. Usually these mistakes are made by researchers and involve oversights such as not using the patched VMs or accidental change of ACLs. Being able to find the mistakes and take corrective action promptly is essential part of best practices. All major Clouds provide logging abilities and we found following on GCP to be the most relevant for catching user mistakes:

- Logging storage access for IP, type of operation (read object, write object, list bucket), which object was affected and number of bytes transferred.
- Logging compute access for user, event time, operation performed, API used to make the access, resource modified (e.g. VM, disks, firewalls, machine images, networks), network traffic in bytes, including notes about whether traffic was between or within compute zones (North Am, Europe, Asia-Pacific, China), or ingress (to the Cloud) or egress (from the Cloud)

Logging and monitoring is typically not a requirement for non-PHI data but we believe that to feel confident about the security implementation, it is a necessary step. Administrators should perform routine logging and mine the logs, using simple scripts or queries, for unexpected access patterns. Following are recommended monitoring practices:

- Google and other Cloud providers send out security bulletins^12^ with details of vulnerabilities and patches. We recommend that administrators monitor that researchers use OS images with the latest security patches.
- Even for a generally HIPAA compliant Cloud provider, like GCP, beta service offerings are not covered by HIPAA. For IRBs guided studies requiring HIPAA compliance, administrator should either disallow service via quota or monitor for use.
- We recommend that administrators stay informed regarding DUA for especially sensitive projects to make sure there are no inadvertent violations.

Note that logging eventually results in data storage and hence cost. Administrators must keep track of these costs and plan for suitable strategies to manage log data volume.

**User training**: We recommend that users take basic HIPAA training to cover IT security, data access, and restrictions and responsibilities for working with sensitive patient information, irrespective of whether on-premise or Cloud systems are used. If such a training is not mandated by the institute, the administrator should put together a basic IT training program around authentication, authorization and data protection guidelines for transferable media.

**Administrator training**: Cloud platforms have abstracted the underlying system administration requirements via high level APIs and GUI, thus removing the burden of understanding low level system engineering. It is our experience that for an existing system administrator or IT savvy informatics personnel, gaining familiarity on Cloud is relatively straightforward. However, we believe that the administrator will need to dedicate time to stay up-to-date with rapidly evolving system management tools via annual trainings.

## Security and Privacy

Security is necessary but not sufficient for privacy. Security can be broken by large scale transnational hacking efforts, occasional rogue behavior, or accidental leaks. All IT systems manage security by monitoring for and actively reacting to security being compromised. We included monitoring as essential part of a secure Cloud. Being able to react in a timely manner to data leaks may require additional methods such as integration with third party solution providers^13^, or application of other machine learning approaches^14^ to identify security breaches in near realtime.

Privacy researchers have shown time and again^15^ that availability of de-identified partial genomic data can result in patient re-identification. The security working group guidelines of Global Alliance for Genomics & Health^16^ (GA4GH) suggests an architecture in which computations themselves are considered to be governed objects within the security framework. Several studies suggest that algorithmic methods such as partial homomorphic encryption^15^, secure multi-party computation^15^ or differential privacy^15^ can provide the necessary privacy protecting layer within such an architecture. The extent to which these algorithmic methods can be integrated with bioinformatics pipelines, queries, statistical and machine learning tools needs further investigation. We believe that such a platform will encourage freer flow of data across silos and reduce complexities around patient consent and DUAs.

